# TME induced TRAIL secretion from engineered macrophages for anti-tumor therapy

**DOI:** 10.1101/2022.04.30.490137

**Authors:** Xin Huang, Botian Deng, Hui Zhou, Binhe Shi, Junhua Liu, Xiaojiao Shan, Xiaobin Fang, Xiushan Yin, Luo Zhang

**Affiliations:** Roc Rock Biotechnology (Shenzhen) Co., Ltd. Shenzhen 518118, China; Department of General Practice Medicine, Shengjing Hospital of China Medical University, Shenyang 110022, China; Research Center of Bioengineering, the Medical Innovation Research Division of Chinese PLA General Hospital, Beijing 100039, China

**Author notes:** Corresponding author: Luo Zhang). Co-corresponding author: Xiushan Yin. These authors contributed to this work equally (Xin Huang, Botian Deng).

## Abstract

Capacity of tumor niche chemotaxis and long-term persistency making macrophage as great vehicles for anti-tumor factor delivery. Macrophages-based delivery of chemokines or cytokines have been tested for tissue homeostasis and cancer repression. TRAIL as a promising anti-tumor cytokine, made little clinical progress due to limited stability and off-target toxicity. Here we engineered macrophages with tumor micro-environment (TME)-induced trimerized CP1-TRAIL secretion under the TME specific promoter Arg1 (Tri-TRAIL-iM). The Tri-TRAIL-iM cells displayed high specific inducible activity in both cell-based co-culture assay and in tumor baring mice models. Compared to normal TRAIL over-expressed macrophages under none-inducible promoter, Tri-TRAIL-iM infiltrated to tumor sites and showed superior apoptosis induction of cancer cells and tumor growth repression as well as less systemic side effect. This inducible delivery TRAIL strategy can be effectively further applied in clinical studies and can be coupled with other engineered methods to maximize the therapeutic outcomes for solid tumors.

## Introduction

Tumor necrosis factor related apoptosis inducing ligand (TRAIL) is a type II transmembrane protein, which is one of several members of tumor necrosis factor superfamily[1, 2]. The main function of TRAIL is to induce apoptosis in transformed cells with high specificity[3]. TRAIL induces apoptosis mainly through the involvement two death receptors (TRAIL-R1/ DR4 and TRAIL-R2/ DR5)[1, 4] in humans and one receptor in mice (mTRAIL-R)[5], and it also induces non apoptotic signaling, including activating nuclear factors-κB (NF-κB), p38, ERK, SRC and Rac1[6]. TRAIL and its receptors are ideal candidates for targeting cancer via extrinsic apoptosis pathway, different versions of TRAIL and TRAIL receptor agonists were developed for clinical application. At present, developing viable treatment candidates such as agonist monoclonal antibodies[7](mapatumumab (HGS-ETR1), lexatumumab (HGS-ETR2), drozitumab (PRO95780), conatumumab (AMG-655), tigatuzumab (CS-1008) and LBY-135)) have been proven challenging, for these agonist mAbs showed little antitumor response in patients. And even in the preclinical tumor xenotransplantation model in mice, mapatumumab and lexatumumab failed to sustainably shrink the tumor, proving that they can only temporarily delay tumor growth[8]. In recent years, a growing number of drug delivery systems (DDS) have been studied to deliver chemotherapeutic and gene therapeutic agents for tumor therapy[9]. However, immunogenicity and cytotoxicity of these DDS to normal tissues and fast-drug metabolism remain as major obstacles to antitumor efficacy, and this hinders their clinical application[10]. Therefore, endogenous DDS including cell-based systems has attracted extensive attention[11, 12].

Macrophages are highly plasticized cells with multiple functions, including tissue development and homeostasis, removal of cell debris, elimination of pathogens and regulation of inflammatory response[13].Cytokines produced by tumor cells and tumor associated stromal cells, such as monocyte chemoattractant protein 1 (MCP-1, also known as CCL-2), macrophage colony-stimulating factor (M-CSF) can recruit monocytes/ macrophages into the tumor microenvironment[14, 15].Macrophage carriers are attractive candidates for drug delivery to tumors because of their affinity to the tumor site[16, 17]. Various macrophage mediated cell carriers have been designed for the treatment of tumor metastasis and solid tumors[17-19]. However, previous studies have mostly used continuous expression of cellular delivery, resulting in non-tumor tissue damage and systemic side effects, making this strategy in need of further improvement.

In this study, we engineered macrophages that specifically express and secrete the cancer cytotoxic protein TRAIL in the context of tumor microenvironment. These engineered macrophages were tested in vitro and in vivo for tumor elimination.

## Methods

### Cell culture

Raw264.7 cells were originally obtained from the American Type Culture Collection (ATCC, Manassas, VA, USA). And were cultured in DMEM medium (High quality 10 % fetal bovine serum (FBS); 0.05 mM β-Mercaptoethanol; 1 % penicillin/streptomycin). Mouse breast cancer cells lines (4T1) were cultured in modified medium DMEM, supplemented with 10 % FBS, 1 % penicillin streptomycin and 1 % fungizone (Invitrogen), and grew in a humid atmosphere of 5 % CO_2_ air at 37 °C until they reached 70-80 % confluence.

### Vector constructs and lentivirus transfection

The lentiviral constructs expressing mouse TRAIL were cloned into the pCDH vector (SBI). TRAIL cDNA was obtained from the National Center for Biotechnology Information. Combining the C-terminal of trail cDNA with the essential h C-propeptide of α1 (I) collagen containing 11 glycine repeat triple chemical region to achieve TRAIL trimer was expressed as a secret protein (Tri-TRAIL-cM) and tumor microenvironment induced trail trimer was marked as Tri-TRAIL-iM. Lentiviral vectors were transfected with packing plasmids into 293T cells for 2 days, and the viral particles were used to infect RAW264.7 cells. Selection was carried out by culturing cells in medium containing 2 μg/mL puromycin for 2 days. RAW264.7 cells with TRAIL were named as Mono-TRAIL-cM and Tri-TRAIL-cM, respectively. For Tri-TRAILl-cM, the Arg1 promoter was applied, and non-TRAILcells were marked as empty-cM.

### Macrophages conditioned medium preparation and co-cultured treatment

To obtain conditioned medium (CM) of RAW264.7, the cell-culture media after 6, 12, and 24 h were collected and centrifuged at 3000 rpm for 10 min at 4 °C. To analyze the effects of macrophages of different conditions on 4T1 cells, the 4T1 cells were then washed and cultured with fresh medium or CM from macrophages for 24 h.

### Cell counting Kit-8 assay (CCK-8)

4T1 cells were co-cultured with different condition medium of macrophages containing various version of TRAIL and controls for 24 h; 4T1 (2×10^3^) were seeded into 96-well plates. Then, 10 μl of CCK-8 (Beyotime, C0037, Jiangsu, China) solution was added to each well at the appointed time. After 1 h of incubation at 37 °C, the absorbance at 450 nm was measured using an automatic microplate reader (Multiskan Spectrum, Thermo Scientific, Waltham, Massachusetts, USA).

### Western blot (WB)

Tissue proteins were extracted according to kit instructions (Sigma Aldrich, R0278, St. Louis, Missouri, USA) and protein concentrations were routinely quantified by using a BCA kit (Beyotime, P0010S, Jiangsu, China). Each sample was analyzed at 30 μg by SDS-PAGE, followed by semi-dry transfer to PVDF membranes. After blocking with 5% skim milk prepared in TBST for 1 h at room temperature, the membranes were incubated overnight at 4 °C with pre-diluted primary antibodies. After that, the membrane was washed and incubated in secondary antibody for 1 hour at room temperature. The dilution ratios of all antibodies were as follows: Myc-Tag (1:500, CST, #2276, Boston, USA), GAPDH (1:500, CST, #97166, Boston, USA).

### Enzyme-linked immunosorbent assay (ELISA)

The levels of murine TRAIL/TNFSF10 (DY1121) in the cells were measured by enzyme-linked immunosorbent assay (ELISA) according to the manufacturer’s instructions (R&D Systems, Emoriville, California, USA). The OD value was measured by Microplate Reader (Multiskan Spectrum, Thermo Scientific, Waltham, Massachusetts, USA).

### Flow cytometry

The surface markers of stimulated cells were detected by flow cytometry. Briefly, RAW264.7 cells were collected from 6-well plates after stimulation for 24 h and washed three times with PBS. The cell suspension respectively containing 1×10^6^ M1-polarized or M2-polarized cells was then divided into 1.5 ml EP tubes and incubated with the following antibodies (APC anti-mouse CD206 (PA5-46879, Thermo Fisher Scientific, USA) on ice for 30 min in the dark. After washed twice, cells were suspended in 500 μl PBS with 3 % FBS and then detected by flow cytometry (BD Biosciences, San Diego, CA, USA).

### Quantitative real-time PCR analysis (qRT-PCR)

Total RNA was extracted from cells by TRIzol and an RNA extraction kit (MiniBEST Universal RNA Extraction Kit, RR820A, Takara, China). Total RNA (1 μg) was reverse transcribed by a reverse transcription kit (RR047A, Takara, China). Real-time quantitative PCR was performed on ABI 7500 system (Applied Biosystems, Foster City, CA) and analyzed using ABI 7500 Software Version 2.03 (Applied Biosystems) according to the manufacturer’s instructions. Relative expression levels of mRNA the target genes were obtained by normalizing to β-actin in each sample and comparing the threshold cycle (CT) values with the calibration curve.

**Table1.**
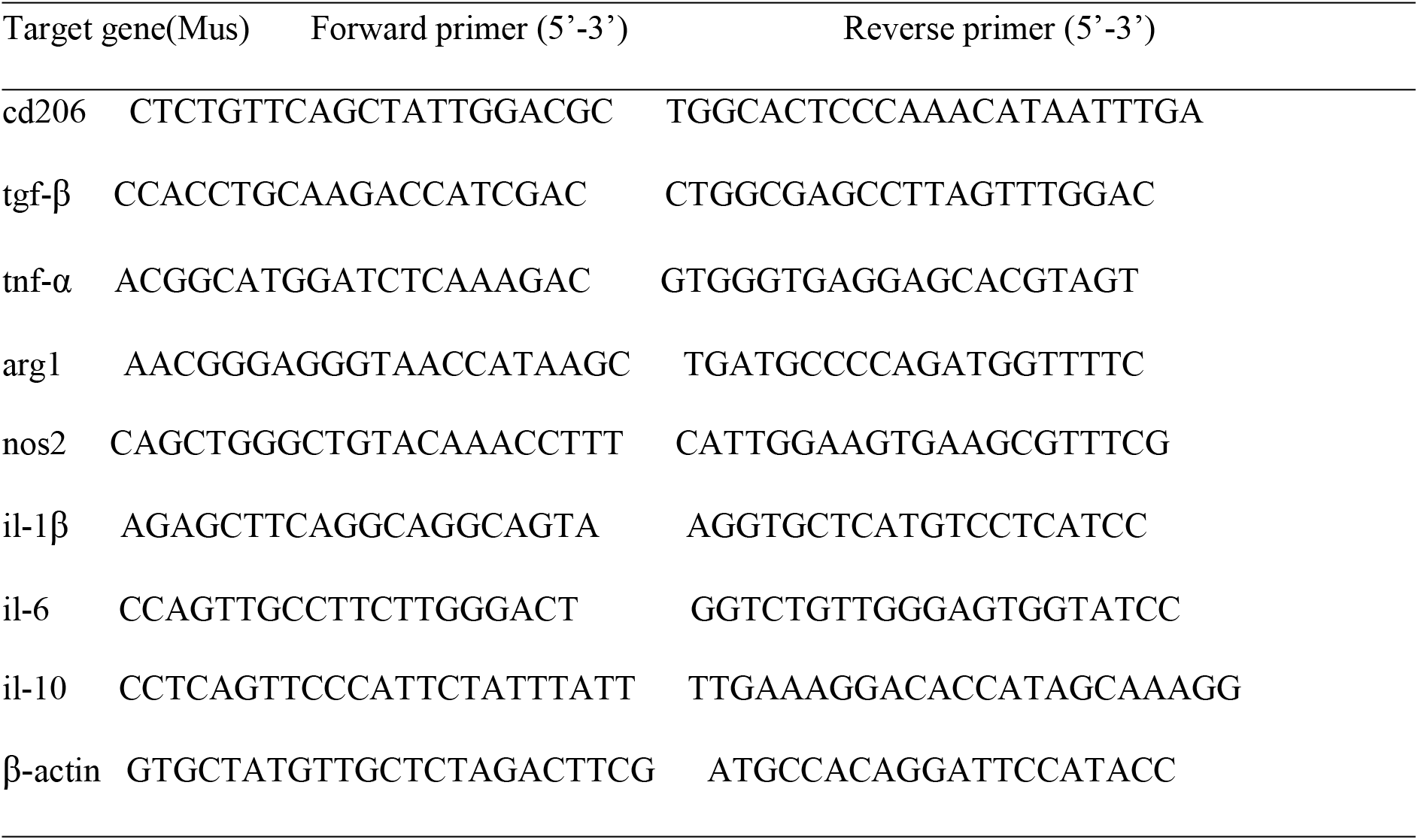
Primer sequences used in real-time PCR analysis.

### Apoptosis Detection by Annexin V/PI Double Staining

Cell apoptosis rate was measured with the Annexin V-FITC Apoptosis Detection Kit (560931, BD, United States) according to the manufacturer’s instructions. Briefly, the RAW264.7 cells were seeded in 6-well plates (5×10^5^/well) then harvested and resuspended in the binding buffer, and co-incubated with 5 μl of Annexin V-FITC and 5 μl of PI for 20 min at room temperature in the dark. The cells were analyzed by a flow cytometry (BD Biosciences, San Diego, CA, USA) within 1 h, and the data analyzed with FlowJo 10.4.2 software (BD Biosciences).

### Tumor Models

Six-ten-week-old male BALB/C mice were purchased from Beijing huafukang Biotechnology Co., Ltd. All animal experiments were conducted in strict accordance with the Guide for the Care and Use of Laboratory Animals of Health’s National Organizations, and all operations performed on mice were approved by the China Medical University Standards for Laboratory Animals Welfare and Ethical Review (Permit Number: KT2022256). Tumor cells were harvested and single cell suspensions in 100 μl PBS (5×10^5^ 4T1 cells) were implanted subcutaneously in the BALB/c mice (for 4T1 model). In the treatment experimental model, 5 × 10^6^ in 100 μl PBS of different RAW264.7 cells (Mono-TRAIL-cM/Tri-TRAIL-cM and empty-cM) or PBS (100 μl) were injected into the tail vein of BALB/C mice. Primary tumor volumes and mice weights were measured 3 times per week using electronic calipers. The maximum longitudinal diameter (length) and maximum transverse diameter (width) were measured. Tumor volume was estimated by a modified ellipsoid formula: volume= 1/2 (long × width^2^)[20]. The tumor tissues were perfused with phosphate-buffered saline and fixed in 10 % paraformaldehyde (PFA) for histological examination.

### Histopathological analysis

Hematoxylin and eosin (H&E) staining (G1120, Solabao, Beijing, China) was used to assess the histochemistry of the tumor tissues. The tissues from mice were fixed in 10 % paraformaldehyde buffer for 24 h, followed by gradient alcohol dehydration, paraffin embedding. Paraffin sections were cut into 5 μm-thick sections and dried in a 37 °C incubator. After xylene dewaxing, the slides were subjected to H&E staining kits according to the manufacturer’s instructions.

### Immunohistochemical (IHC) and immunofluorescence (IF) staining

The tissue was sectioned 5 μm thick. Serial sections were placed on slides and dried in a 37ºCm incubator. After dewaxing with xylene, the sections were immersed for 2 min in successive 50 %, 70 % and 80 % ethanol, respectively, and in eosin for 5 s. The specimens were then routinely dehydrated, hyalinized, and fixed. The sections were rinsed with water and then subjected to antigenic heat repair, incubated with TRAIL (1:500, Abcam, ab231063), followed by secondary immunoglobulin G coupled with appropriate biotin (1: 1000, Affinity, #S0001) incubation. These sections were stained with hematoxylin for 5 min, rinsed with tap water for 3 min, 1 % hydrochloric acid for 2 s, then tap water for 2 min, before being sealed by gradient dehydration. For IF staining, specimen was covered with fluorescent-labeled antibody (iNOS, 1:500, Abcam, ab178945; F4/80, 1:500, Abcam, ab60343) and stored in an enamel box for 30 min. The secondary-fluorescence immunoglobulin G was from Abmart (1:1000, M21014S, Shanghai, China). The co-stained sections were examined, and images were captured by a Nikon Eclipse 90i fluorescence microscope.

### Tumor immune evaluation

RNA-sequencing expression profiles and corresponding clinical information for breast cancer were downloaded from the TCGA dataset(https://portal.gdc.com). To assess the reliable results of immune score evaluation, we used immuneeconv. It’s an R software package that integrates six latest algorithms, including TIMER, xCell, MCP-counter, CIBERSORT, EPIC and quanTIseq. These algorithms had been benchmarked, each had a unique advantage. The algorithm we adopted in this study was EPIC. The R software ggstatsplot package was used to draw the correlations between gene expression and immune score Used Spearman’s correlation analysis to describe the correlation between quantitative variables without a normal distribution. P values less than 0.05 were considered statistically significant (*P < 0.05). All the above analysis methods and R package were implemented by R foundation for statistical computing (2020) version 4.0.3 and software packages ggplot2 and pheatmap.

### TUNEL assay

DNA fragments were analyzed by TUNEL (transferase dUTP nick end labeling) method. After dewaxing, the sample was rehydrated with graded ethanol and immersed in PBS with 4 % formaldehyde for 20 minutes. The samples were treated with click it plus in situ apoptosis detection, TUNEL analysis and Alexa fluor 488 dye (c10617; Thermo Fisher). The nuclei were stained and installed with vectashield installation medium (h-1400; vector laboratories, California, USA). Ki-67 was used as markers of proliferation with red fluorescence. TUNEL positive cells produce green fluorescence. Immunofluorescence images were obtained using a fluorescence microscope (Olympus).

### Data analysis

All statistical analyses were performed using GraphPad Prism 8.0 (GraphPad Inc, La Jolla, CA, USA). For the measurable data, continuous variables were expressed as mean± standard deviation (x± s). One-way ANOVA was used for comparison between groups. Tukey test was used for pairwise comparison. ANOVA and Mann Whitney U test was used to compare the data among different groups. For all tests, a two-tailed P value of <0.05 was recognized to be statistically significant.

## Results

### Characteristics of Mono-TRAIL-cM and Tri-TRAIL-cM

We engineered RAW264.7 cells that secrete murine native TRAIL (Mono-TRAIL-cM) and trimeric TRAIL (Tri-TRAIL-cM) by combining the human C-propeptide of α1(I) collagen (Trimer-Tag) in-frame fusion with the C-terminus of mature murine TRAIL, leading to a disulfde bond-linked homotrimer respectively (Fig. 1A). Secreted TRAIL proteins were further quantified and the ELISA results showed that TRAIL concentration in the cell supernatant was positively correlated with culture period (Fig. 1B). Next, we investigated the polarization of the engineered macrophagesby detecting the constructed cells by flow cytometry, as evident, and in contrast to the non engineered ones, engineered macrophages reversed the TME induced conversion of macrophages to an anti-inflammatory type (Fig. 1C). Meanwhile, qRT-PCR showed that engineered macrophages showed more transcriptional activity of pro-inflammatory genes, and Tri-TRAIL-cM still showed strong transcriptional activity of pro-inflammatory genes under TME induction. It should be noted that arg1 showed specific transcriptional activity induced by TME (Fig. 1D).

**Figure 1.**
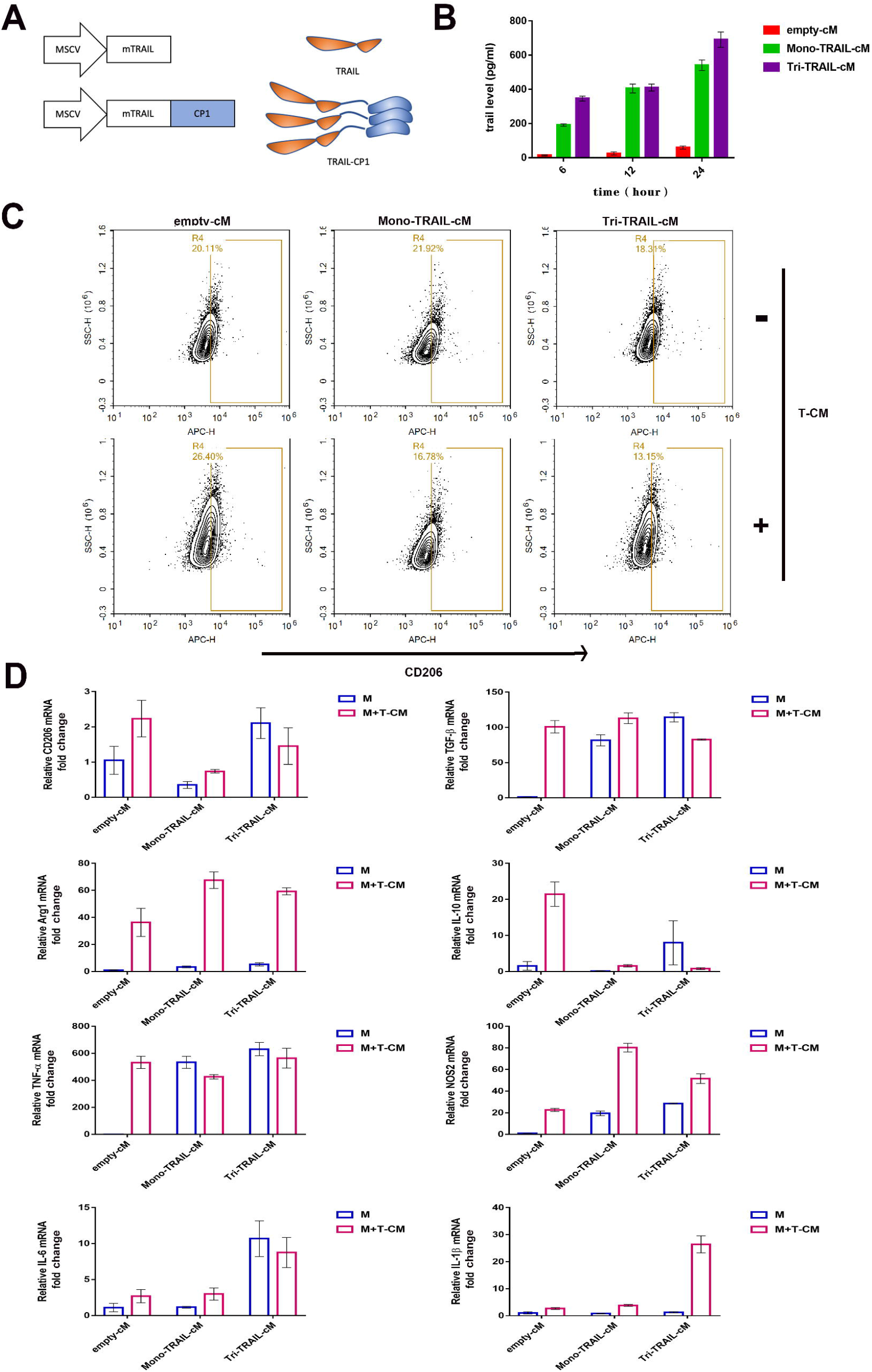
Characterization of secreted macrophage drug carriers. (A) Construction of vector architectures for Mono-TRAIL and Tri-TRAIL, schematic of mono- and tri-architectures for secreted TRAIL. (B) The amount of TRAIL secreted by empty-cM, Mono-TRAIL-cM and Tri-TRAIL-cM in different duration (pg/ml). (C) Flow cytometry showed the expression of CD206 of empty-cM, Mono-TRAIL-cM and Tri-TRAIL-cM under different conditions (T-CM: tumor conditioned medium). (D) The mRNA relative expression of cd206, tgf-β, arg1, il-10, tnf-α, nos2 and il-1β in different cell lines. \**P*<0.05.

### In vitro antitumor effect of Mono-TRAIL-cM and Tri-TRAIL-cM

The antitumor effect of Mono-TRAIL-cM/Tri-TRAIL-cM and empty-cM on the 4T1, B16, CT26 cells was examined by CCK-8 and living cell staining at 6,12 and 24 h (Fig. 2A and B), respectively. Compared with the empty-cM group, the engineered macrophages had better antitumor activity in co-culture system from which the supernatant were collected, except CT26. Among them, we found Tri-TRAIL-cM had the most significant antitumor effect (Fig. 2C and D). The activity of 4T1 cells were inhibited the highest with the Tri-TRAIL-cM supernatant. Mono-TRAIL-cM supernatant had mild antitumor but significant effect compared to control cells. However, in CT26 cells, cytotoxicity was visualized only with the conditioned medium containing of Tri-TRAIL-cM collected at 24 h. The results indicate trimerized TRAIL had superior tumor cell killing ability in vitro. We further analyzed the apoptosis of 4T1 cells after co-culture, and the results showed that Tri-TRAIL-cM had a stronger pro-apoptotic effect (Fig. 2F and G).

**Figure 2.**
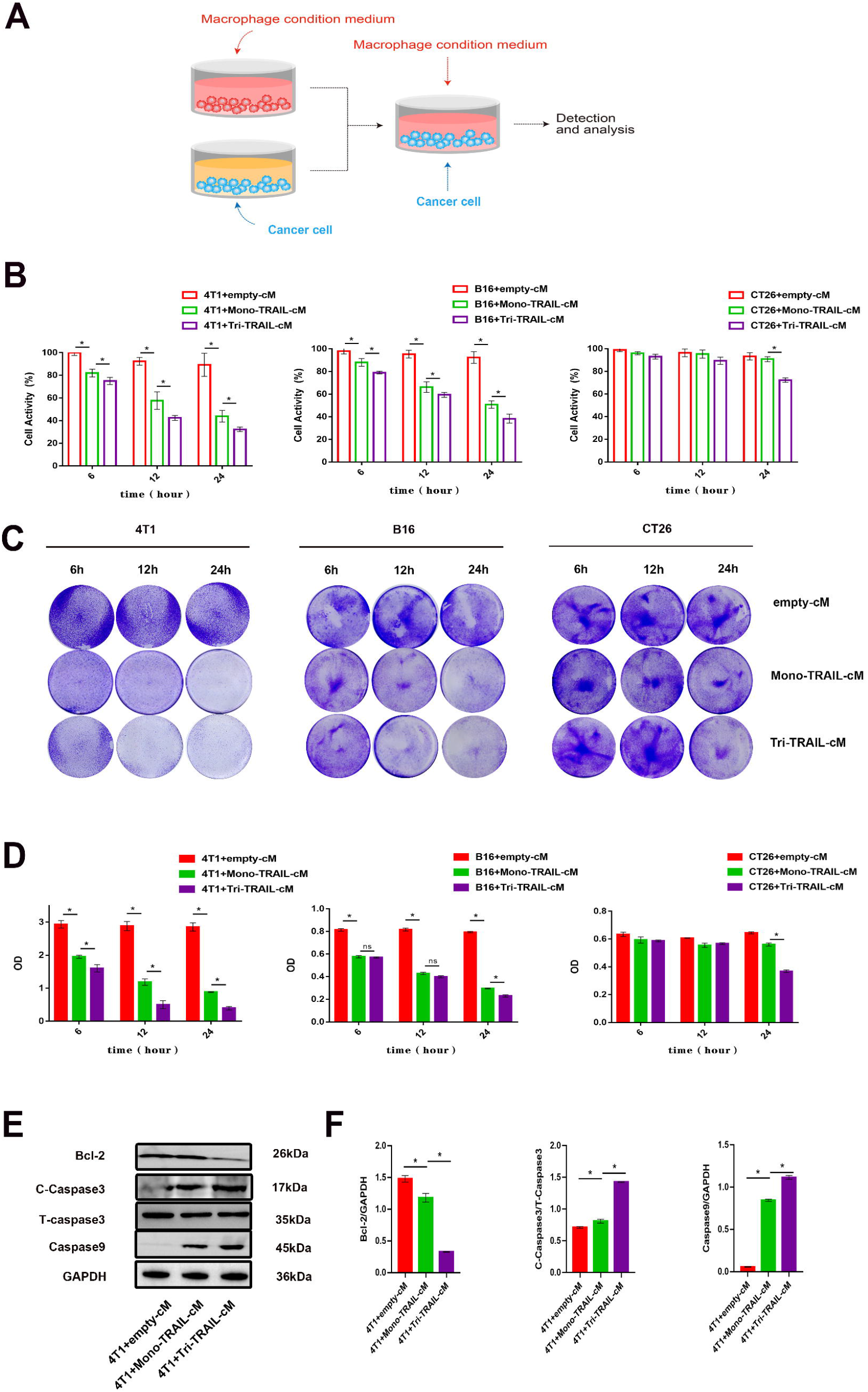
Antitumor activity of secretory macrophage drug carrier in vitro. (A) Schematic diagram of co-culture assay. (B) CCK8 validated the effect of macrophage conditioned media collected at different time points on tumor cell activity. (C) Crystal violet staining of tumor cells cultured with macrophage conditioned media. (D) Quantification of crystal violet staining of cells. (E-F) The protein relative expression of apoptosis related proteins Bcl-2, Cleaved-Caspase 3/Total-Caspase 3 and Caspase 9 of 4T1 after co-culture. \**P*<0.05.

### Characteristics of Tri-TRAIL-iM

To avoid unspecific killing and enhance the antitumor specificity of Tri-TRAIL-cM, we took advantage of inducible promoters that regulated by the tumor microenviroment in vivo. Previous experiments showed Arg1 was highly expressed in RAW264.7 cells, after exposure to tumor conditioned medium (TCM). Arg1 is able to respond to IL-10 and IL-4 highly expressed by the tumor environment, whereas under other inflammatory factor stimulated conditions or in inflammatory models, Arg1 expression of RAW264.7 was largely unaffected [21].

We replaced the constitutively expressed CMV promoter of Tri-TRAIL-cM to the Arg1 promoter/enhancer, and generated the tumor environment induced secretion of macrophage Tri-TRAIL-iM (Fig. 3A and B). Under normal conditions, Tri-TRAIL-iM secreted only low doses of TRAIL, whereas Tri-TRAIL-iM secreted TRAIL protein in large amounts after co-culture with 4T1, (Fig. 3C), indicating the feasible response of Arg1 promoter to tumor associated stimuli in the presence of 4T1. We next observed the induced secretion of TRAIL and the results show that only under the stimulation of 4T1 cells, RAW264.7 cells can highly express TRAIL protein (Fig. 3D). Under normal culture, Tri-TRAIL-iM behaved as a non-inflammatory macrophage phenotype, whereas under TME induction, Tri-TRAIL-iM exhibited transcriptionally active activation of proinflammatory genes (Fig. 3E and F). We further verified the induced expression of Tri-TRAIL-iM in tumor tissues, and the results showed that more TRAIL expression was observed in Tri-TRAIL-iM groups by IHC (Fig. 3G).

**Figure 3.**
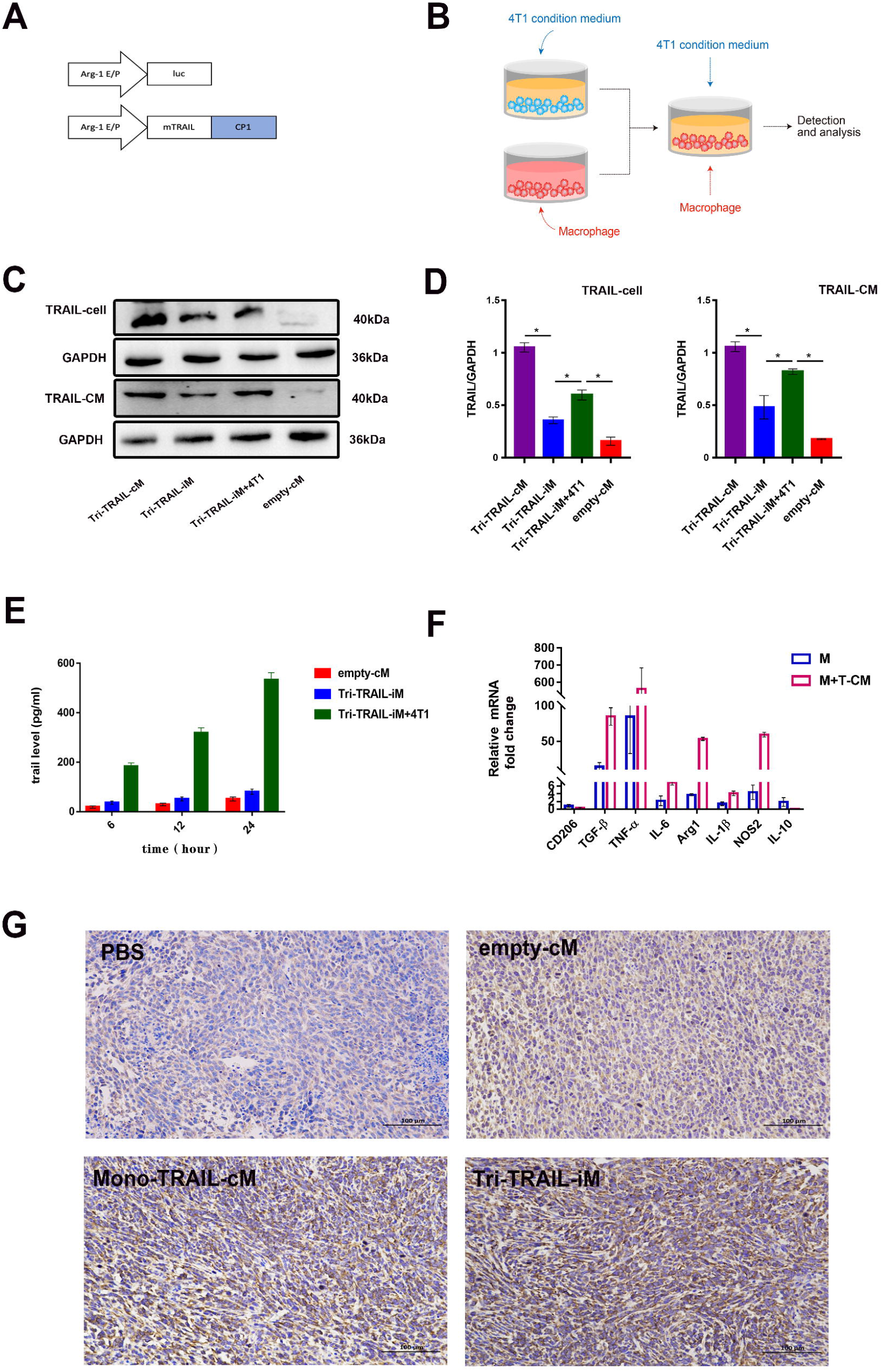
Characterization of Tri-TRAIL-iM. (A) Schematic representation of the vectors for Arg1-luciferase and Arg1-Tri-TRAIL. (B) Diagram of the co-culture assay by which the TME induces tri-TRAIL secretion. (C-D) Protein relative expression of TRAIL in cells and medium (CM) of different groups (\**P*<0.05). (E) The amount of TRAIL protein secretion of Tri-TRAIL-iM induced by TME in different time lengths (pg/ml). (F) The mRNA relative expression of cd206, tgf-β, arg1, il-10, tnf-α, nos2 and il-1β of Tri-TRAIL-iM (M: Tri-TRAIL-iM; T-CM: the conditioned medium of 4T1 cells). (G) Immuno-staining of TRAIL in tumor tissues of 4T1 tumor bearing mice.

### In Vivo anti-tumor effect of Tri-TRAIL-iM in a mouse tumor model

Next the antitumor effect of Tri-TRAIL-iM was evaluated in subcutaneous tumor bearing mice (Fig. 4A). We measured the tumor volumes in the PBS, empty-cM, Mono-TRAIL-cM, and Tri-TRAIL-iM groups at the days as indicated, respectively. For individual tumor growth (Fig. 4B), we found a burst growth of tumors in none of the mice in the Tri-TRAIL-iM treated group compared to the other groups, with a trend of tumor growth for all mice within the groups. For the overall tumor volume (Fig. 4C), there was no significant difference in the tumor volume of the PBS group and the empty-cM group, indicating the limited antitumor effects of none TRAIL RAW264.7 cells. Mono-TRAIL-cM had weak killing effect on the tumor and the Tri-TRAIL-iM was able to effectively reduce the tumor volume. At the same time, we observed that the trend of weight growth treated with PBS, empty-cM and Mono-TRAIL-cM slowed down and even there was a drop in weight at the later stages of the experiment, while the overall weight of Tri-TRAIL-iM treated tumor bearing mice kept increasing (Fig. 4D). According to the TUNEL assay results, we found compared with the PBS, empty-cM groups, the tumor showed stronger apoptosis signals and weaker proliferation ability, the number of TUNEL positive cells significantly increased and the expression of Ki-67, a cellular marker of proliferation, was significantly suppressed in the tumors (Fig. 4E). Besides, there were more pro-inflammatory macrophages (expressed the iNOS and F4/80) in Mono-TRAIL-cM, and Tri-TRAIL-iM groups, respectively (Fig. 4F).

**Figure 4.**
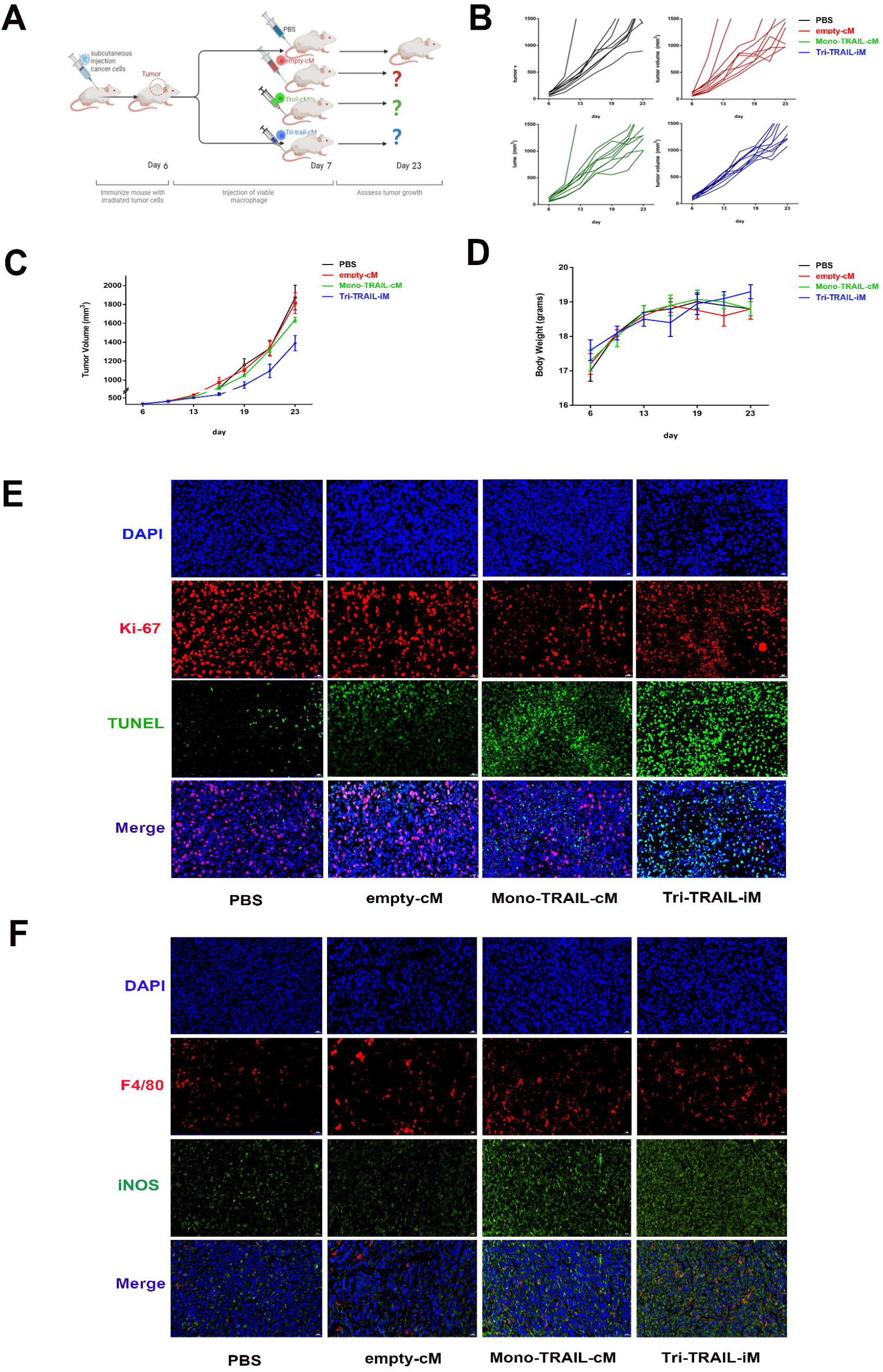
Antitumor activity of Tri-TRAIL-iM in vivo. (A) Diagram of Tri-TRAIL-iM treatment in tumor bearing mice. (B) Individual tumor volume changes in tumor bearing mice with different treatment. (C) Overall tumor volume changes in tumor bearing mice with different treatment. (D) Body weight changes of tumor bearing mice with different treatment. (E) Ki-67 (red fluorescence) and TUNEL (green fluorescence) in tumor tissues of mice. (F) Immunofluorescence co-localization by F4/80 (green fluorescence) and iNOS (red fluorescence) in mouse tumor tissues.

### TRAIL expression correlates with immune effector cells and uncharacterized cells in breast cancer

To investigate the relevance of our observations in human cancer, we explored the correlation of macrophage infiltration and TRAIL expression with breast cancer in the Cancer Genome Atlas (TCGA) dataset. We found that in breast cancer patient tumor tissues, macrophage associated gene expression was relatively conserved compared with that in non tumor tissues (Fig. 5A), predicting that the macrophage occupancy of tumor tissues was relatively stable and that as a drug carrier macrophages themselves might be less able to alter the immune environment of breast cancer tumor tissues. At the same time we explored the potential impact of TRAIL on the immune landscape in breast cancer, and we found a positive correlation between TRAIL transcripts and a gene signature specific to human CD4 effector T cells in cancer patients and a negative correlation with immune signature free cells (Fig. 5C). We reasoned that TRAIL might be important for the responsiveness to therapies that promote CD4 responses. The negative correlation with the most represented uncharacterized cells in breast cancer indicated besides the anti-tumor activity, TRAIL might be an additional biomarker for tumor outcome (Fig. 5B).

**Figure 5.**
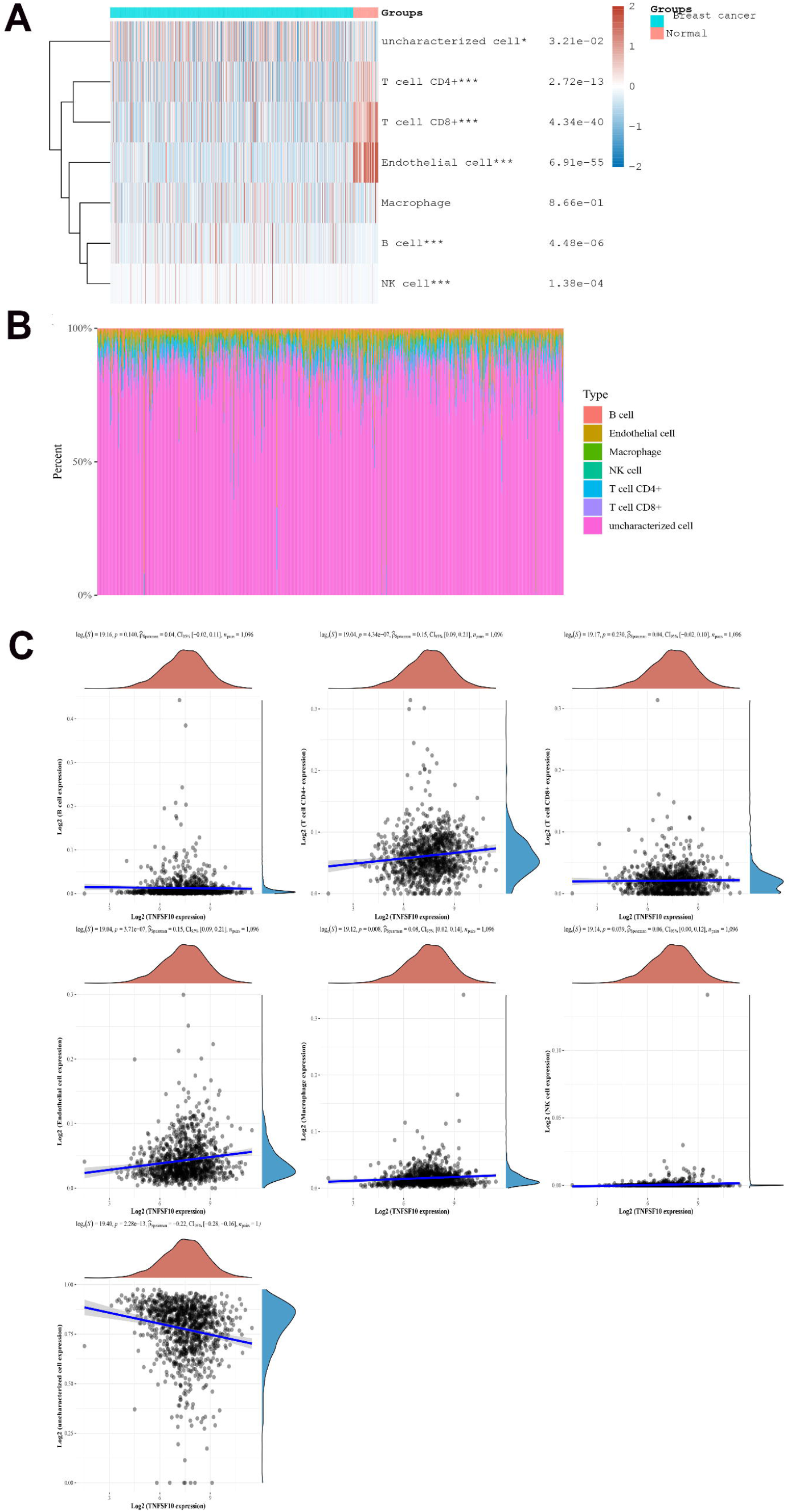
Macrophage related transcriptome signature and TRAIL expression in breast cancer. (A) Immune cell score heatmap, different colors represented different expression distribution in different samples. The statistical difference of two groups was compared through the Wilcox test. (B) The percentage abundance of tumor infiltrating immune cells in each sample. Different colors represents different types of immune cells. The abscissa represents the sample, and the ordinate represents the percentage of immune cell content in a single sample. (C) The correlations between gene expression and immune score was analysed with Spearman.The abscissa represents the distribution of the gene expression or the score, and the ordinate represents the distribution of the immune score. The density curve on the right represents the trend in distribution of the immune score, the upper density curve represents the trend in distribution of the gene expression or the score. The value on the top represents the correlation p value, correlation coefficient and correlation calculation method. \**p* < 0.05, \*\**p* < 0.01,\*\*\**p* < 0.001.

## Discussion

Due to the tropism for hypoxia and the ability to migrate and infiltrate into solid tumors, myeloid lineage has been explored as potential carriers for anticancer drugs and imaging agents[22, 23]. Based on this we designed a drug carrier for macrophages capable of secreting TRAIL, one of the TNF superfamily members able to exert a pro-apoptotic function towards TRAIL sensitive cancer cells and spar normal cells. The macrophages were armed with Mono and Tri-TRAILs and the antitumor activity of modified macrophages were compared in vitro and in vivo. Trimerized TRAIL by CP1 peptide linker showed enhanced cytotocix activity, which is consistent with the observation that trimerization is necessary to generated a more efficient apoptosis[24].

Macrophages are the major immune cells in most solid tumors[21]. In mice, tumor associated M2 polarization is characterized by the upregulation of genes involved in fostering an immunosuppressive microenvironment[25]. The tumor microenvironment (TME) is characterized by acidosis, hypoxia, elevated concentrations of IL-4 and IL-13 and tumor derived cytokines[26].A time-dependent increase in the expression of the M2 markers Ym1, FIZZ1, Mrc1 and Arg1 has been reported[21]. After exposure to tumor conditioned medium (TCM), BMDMs and RAW264.7 macrophages also upregulate Arg1 up to 600 folds [21]. This effect may be mediated by tumor derived cytokines and metabolic intermediates, including lactate. Based on this, we constructed the Tri-TRAIL-iM that initiates secretion of Tri-Trail under Arg1 promotor, taking advantage of the tumor reprogrammed macrophages with high Arg1 induction. TRAIL secretion by Tri-TRAIL-iM was proportional to TCM culture time, and the secretion of TRAIL remained at a very low baseline level in the none-induced condition (Fig. 3).

In mice, Tri-TRAIL-iM correlated with high amounts of secreted TRAIL mainly at the tumor site, in contrast to the situation in healthy mice where TRAIL secretion was barely detectable. Meanwhile, in tumor bearing mice, Tri-TRAIL-iM showed a better antitumor effect. These results suggest that Tri-TRAIL-iM has a better safety profile while combating tumors (Fig 4). Moreover, owing to the high expression of TRAIL locally in tumors, which further converts TAMs into pro-inflammatory cells, our strategy could antagonize the pro-tumor activity of naturally occurring tumor infiltrating cell types (Fig 6 B).

In summary, this study demonstrated that Tri-TRAIL has a pro apoptotic effect to control solid tumor growth when delivered locally into tumor tissue by long-lived macrophages, and thus this tumor environment induced drug secreting cells that might merit further investigation as potential candidates for precise cell configuration that could be combined with therapies such as CAR-engineered cells, providing a better efficacy profile to diseases other than cancer.

## Ethics approval and consent to participate

The study was approved by the Ethics Committee of China Medical University (KT2022256).

## Consent for publication

Not applicable

## Availability of data and materials

The datasets are available from the corresponding author on reasonable request.

## Authors’ contributions

Xiushan Yin and LuoZhang designed and supported experiments. Xin Huang and Botian Deng performed the experiments, analyzed data and wrote the manuscript. Binhe Shi and Junhua Liu performed the experiments and assisted mouse breeding and analysis of data. Hui Zhou, Xiaojiao Shan performed experiments. All authors read and approved the final manuscript.

## Acknowledgements

Not applicable

